# Unbiased Precision Estimation under Separate Sampling

**DOI:** 10.1101/342626

**Authors:** Shuilian Xie, Ulisses M. Braga-Neto

## Abstract

**Motivation:** Precision and recall have become very popular classification accuracy metrics in the statistical learning literature. These metrics are ordinarily defined under the assumption that the data are sampled randomly from the mixture of the populations. However, observational case-control studies for biomarker discovery often collect data that are sampled separately from the case and control populations, particularly in the case of rare diseases. This discrepancy may introduce severe bias in classifier accuracy estimation.

**Results:** We demonstrate, using both analytical and numerical methods, that classifier precision estimates can display strong bias under separating sampling, with the bias magnitude depending on the difference between the case prevalences in the data and in the actual population. We show that this bias is systematic in the sense that it cannot be reduced by increasing sample size. If information about the true case prevalence is available from public health records, then a modified precision estimator is proposed that displays smaller bias, which can in fact be reduced to zero as sample size increases under regularity conditions on the classification algorithm. The accuracy of the theoretical analysis and the performance of the proposed precision estimator under separate sampling are investigated using synthetic and real data from observational case-control studies. The results confirmed that the proposed precision estimator indeed becomes unbiased as sample size increases, while the ordinary precision estimator may display large bias, particularly in the case of rare diseases.

**Availability:** Extra plots are available as Supplementary Materials.

**Author summary:** Biomedical data are often sampled separately from the case and control populations, particularly in the case of rare diseases. Precision is a popular classification accuracy metric in the statistical learning literature, which implicitly assumes that the data are sampled randomly from the mixture of the populations. In this paper we study the bias of precision under separate sampling using theoretical and numerical methods. We also propose a precision estimator for separate sampling in the case when the prevalence is known from public health records. The results confirmed that the proposed precision estimator becomes unbiased as sample size increases, while the ordinary precision estimator may display large bias, particularly in the case of rare diseases. In the absence of any knowledge about disease prevalence, precision estimates should be avoided under separate sampling.

## 1 Introduction

Biomarker discovery is typically attempted by means of observational case-control studies where classification techniques are applied to high-throughput measurement technologies, such as DNA microarrays [1], next-generation RNA sequencing (RNAseq) [2], or “shotgun” mass spectrometry [3]. The validity and reproducibility of the results depend critically on the availability of accurate and unbiased assessment of classification accuracy [4, 5].

The vast majority of published methods in the statistical learning literature make the assumption, explicitly or implicitly, that the data for training and accuracy assessment are sampled randomly, or unrestrictedly, from the mixture of the populations. However, observational case-control studies in biomedicine typically proceed by collecting data that are sampled with restrictions. The most common restriction, and the one that is studied in this paper, is that the data are sampled separately from the case and control populations. This is always true in studies involving rare diseases, since sampling randomly from the population at large would not yield enough cases. That creates an important issue in the application of traditional statistical learning techniques to biomedical data, because there is no meaningful estimator of case prevalences under separate sampling. Therefore, any methodology that directly or indirectly uses estimates of case prevalence will be severely biased.

*Precision* and *Recall* have become very popular classification accuracy metrics in the statistical learning literature [6–8]. In this paper, we investigate the bias of precision and recall sample estimates when the typical separate sampling design used in case-control studies is not properly taken into account. Synthetic and real-world biomedical data are used to quantify the magnitude of the bias, which is systematic in the sense that it cannot be reduced by increasing sample size. If information about the true prevalence of cases is available, then a modified estimator is proposed that displays smaller bias, which can be decreased to zero asymptotically as sample size increases under certain regularity conditions on the classification algorithm, in a sense to be made precise.

In [9], a similar study was conducted into the accuracy of cross-validation under separate sampling. It was shown that the usual “unbiasedness” property of k-fold cross-validation does not hold under separate sampling. In fact, the bias can in fact be substantial and systematic, i.e., not reducible under increasing sample size. In [9], modified k-fold cross-validation estimators were proposed for the class-specific error rates. In the case where the true case prevalence is known, those estimators can be combined into an estimator of the overall error rate, which satisfies the usual “unbiasedness” property of cross-validation.

The present paper employs analytical and numerical methods to show that the ordinary precision estimator can display large bias under separate sampling. More specifically, while the recall estimator is asymptotically unbiased as sample size increases, under regularity conditions on the classification rule to be specified, the precision estimator may display a systematic bias, which cannot be reduced by increasing sample size if the observed prevalence of cases in the data is different from the true prevalence in the population of interest. This is a consequence of the fact that precision is a function of the prevalence, whereas recall is not. Case-control studies involving rare diseases are specially affected, since in those studies the true prevalence is small and will almost always differs substantially from the observed prevalence in the data. To address this problem, we propose a new estimator for precision, which can be applied in case the true prevalence is known. This estimator has small bias that vanishes as sample size increases under certain regularity conditions. In the absence of any knowledge about the prevalence, precision estimates should be avoided under separate sampling.

## 2 Materials and Methods

In this section we define and investigate the various error rates of interest in this study, including precision and recall.

### 2.1 Population Performance Metrics

The *feature* vector **X** ∊ *R^d^* summarizes numerical characteristics of a patient (e.g, blood concentrations of given proteins). The *label Y* ∊ {0,1} is defined as *Y* = 0 if the patient is from the control population, and *Y* = 1 if the patient is from the case population. The *prevalence* is defined by

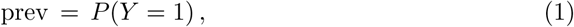

i.e., the probability that a randomly selected individual is a case subject. The prevalence plays a fundamental role in the sequel.

A *classifier ψ*: *R^d^* → {0,1} assigns **X** to the control or case population, according to whether *ψ*(**X**) = 0 or *ψ*(**X**) = 1, respectively. The classification sensitivity and specificity are defined as:

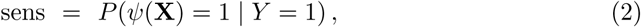

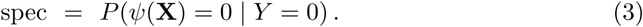

The closer both are to 1, the more accurate the classifier is. A noteworthy property of the sensitivity and specificity is that they *do not depend on the prevalence.*

Other common performance metrics for a classifier are the *false-positive* (FP), *false-negative* (FN), *true-positive* (FP), and *true-negative* (FN) rates, given by

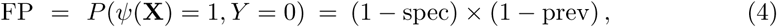

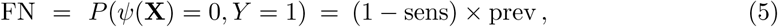

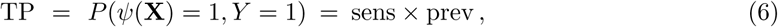

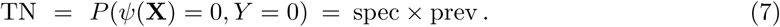

Unlike sensitivity and specificity, the previous performance metrics *do* depend on the prevalence. See Fig. 1 for an illustration.

**Fig 1.**
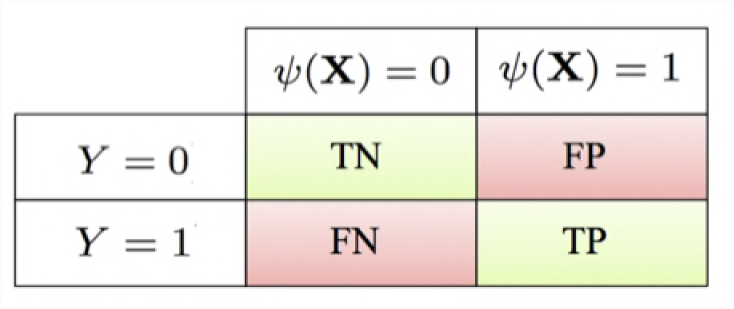
Diagram of error (red) and accuracy (green) rates.

Notice that

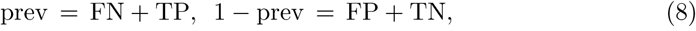

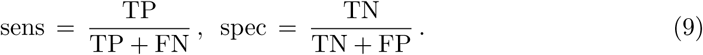

Finally, we define the precision and recall accuracy metrics. Precision measures the likelihood that one has a true case given that the classifier outputs a case:

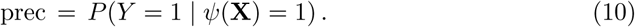

Applying Bayes’ Theorem and using previously-derived relationships reveal that:

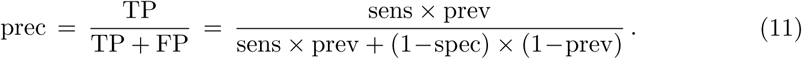

On the other hand, recall is simply the sensitivity:

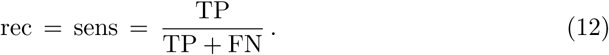

It follows that precision depends on the prevalence, but recall does not.

### 2.2 Estimated Performance Metrics

In practice, the performance metrics defined in the previous section need to be estimated from sample data *S_n_* = {(X_1_, *Y*_1_), …, **X***_n_*, *Y_n_*)}. Let *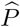* denote the empirical probability measure defined by *S_n_.* The estimator of prevalence is:

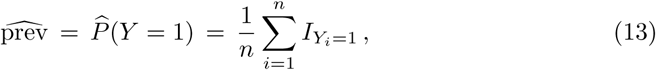

where *I_A_* = 1 if *A* is true and *I_A_* = 0 if *A* is false. Similarly,

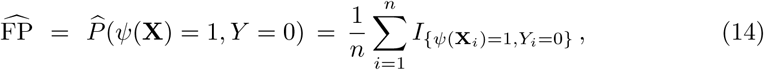

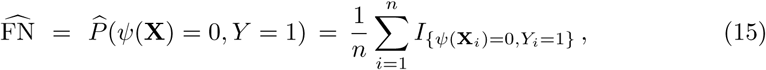

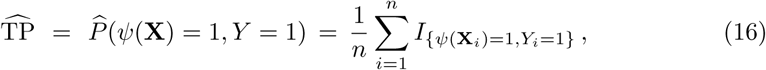

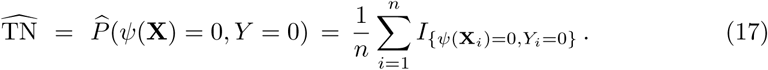

The remaining performance metrics estimators are defined analogously, using (9), (11), and (12):

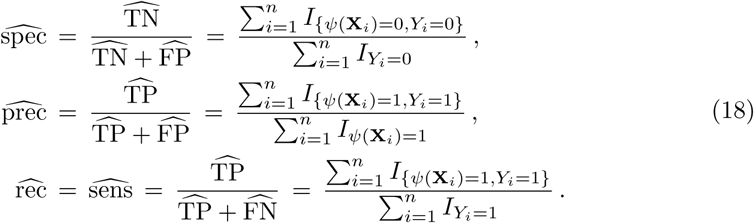

### 2.3 Mixture and Separate Sampling

The usual scenario in Statistical Learning is to assume that *S_n_* = {(**X**_1_, *Y*_1_), …, **(X***_n_*, *Y_n_*)} is an independent and identically distributed (i.i.d.) sample from the true distribution of the pair **(X,** *Y*).That makes *S_n_* a sample from the *mixture* of populations, where each label *Y_i_* is distributed as:

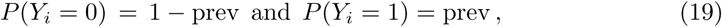

for *i* = 1, …, *n.* Under mixture sampling, 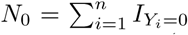 and 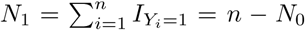 are binomial random variables, with parameters (*n*, 1 − prev) and (*n*, prev), respectively.

By contrast, observational case-control studies in biomedicine typically proceed by collecting data from the populations separately, where the separate sample sizes *n*_0_ and *n*_1_, with *n*_0_ + *n*_1_ = *n*, are pre-determined and nonrandom; i.e., sampling occurs with the restriction *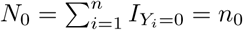* (or, equivalently, *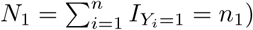*). Therefore, all probabilities and expectations over the sample are conditional on *N*_0_ = *n*_0_. The restriction means that the labels *Y*_1_, …, *Y_n_* are no longer independent, even though the feature vectors **X**_1_, …, **X***_n_* are still independent given the labels. It is not difficult to verify that

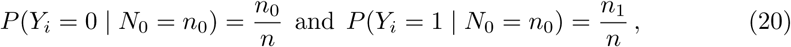

for *i* = 1, …, *n*. Comparing (19) and (20) reveals the main difference between mixture and separate sampling.

### 2.4 Bias of Precision and Recall Estimators

In this subsection, we present an approximate large-sample analysis of the bias of the estimators discussed previously, focusing on the precision and recall estimators. Estimation bias is defined as the expectation over the sample data *S_n_* of the difference between the estimated and true quantities.

The situation is clear with the estimator of the prevalence itself, given by (13). Under mixture sampling, we have

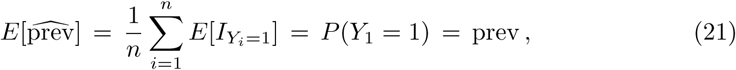

so the estimator is unbiased (in addition, as *n* increases, 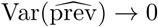 and 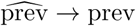 in probability, by the law of large numbers). However, under separate sampling,

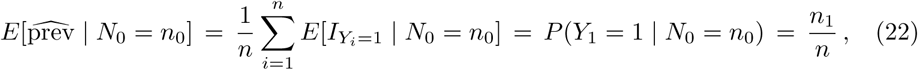

according to (20). This also follows directly from the fact that 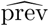 becomes a constant estimator, 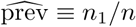, according to (13). Thus,

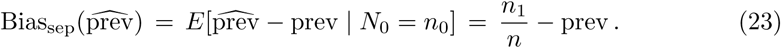

Assuming that the ratio *n*_1_*/n* is held constant as *n* increases (e.g., the common balanced design case, *n*_0_ = *n*_1_ = *n*/2), then this bias cannot be reduced with increased sample size. Furthermore, the bias is larger the further away the true prevalence is from 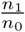. In particular, the bias will be large when prev is small, as in case-control studies involving rare diseases.

The situation for 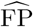, 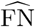, 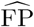, and 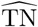 is more complicated. First, we are interested in a classifier *ψ_n_* derived by a classification rule from from the sample data *S_n_* = {(**X**_1_, *Y*_1_), …, **X***_n_, Y_n_*)}. Therefore, all expectations and probabilities in the previous sections are conditional on *S_n_.* Under mixture sampling, the powerful *Vapnik Chervonenkis Theorem* can be applied to show that all of these estimators are asymptotically unbiased, provided that classification rule has a finite *VC Dimension* [10]. This includes many useful classification algorithms such as LDA, linear SVMs, perceptrons, polynomial-kernel classifiers, certain decision trees and neural networks, but it excludes nearest-neighbor classifiers, for example. Classification rules with finite VC dimension do not cut the feature space in complex ways and are thus generally robust against overfitting.

Assuming mixture sampling and a classification algorithm with finite VC dimension *V_C_*, it can be shown that (details omitted; see [5, p. 47] for a similar argument)

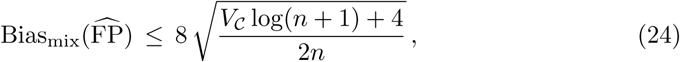

so that the bias vanishes as *n* → ∞. Similar inequalities apply to 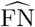, 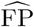, and 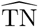. These are distribution-free results, hence vanishingly small bias is guaranteed if *n* ≫ *V_C_,* regardless of the feature-label distribution. For linear classification rules, *V_C_* = *d* + 1, where *d* is the dimensionality of the feature vector. In this case, the 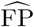, 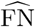, 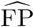, and 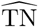 estimators are essentially unbiased if *n* ≫ *d*.

Next we consider the bias of the precision and recall estimators under mixture sampling (the analysis for the sensitivity and specificity estimators is similar; in fact, the latter is just the recall estimator). We will make use of the following relation for the expectation of a ratio of two random variables *W* and *Z*:

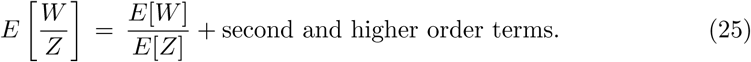

This equation can be proved by expanding *W/Z* around the point (*E*[*W*], *E*[*Z*]) using a bivariate Taylor series and taking expectation (details are omitted for space). The approximation obtained by dropping the higher order terms is quite accurate if *W* and *Z* are around *E*[*W*] and *E*[*Z*], respectively (it is asymptotically exact as *W* → *E*[*W*] and *Z* → *E*[*Z*]). For the precision estimator,

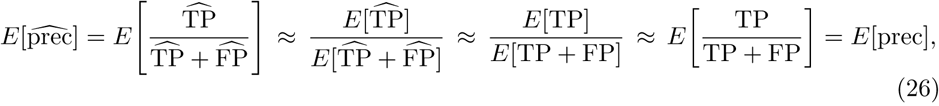

for a sufficiently large sample, where we used the previously-established asymptotic unbiasedness of 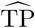, 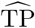, and 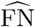. An entirely similar derivation shows that 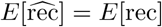. Hence, for “well-behaved” classification algorithms (those with finite VC dimension), both the precision and recall estimators are asymptotically unbiased under mixture sampling.

Unfortunately, there is not at this time a version of VC theory for separate sampling. In order to obtain approximate results for the separate sampling case, we will assume instead that, at large enough sample sizes, the classifier *ψ* is nearly constant, and invariant to the sample. This assumption is not unrelated to the finite VC dimension assumption made in the case of mixture sampling. Many of the same classification algorithms that have finite VC dimension, such as LDA and linear SVMs, will also become nearly constant as sample size increases. In this case, we have

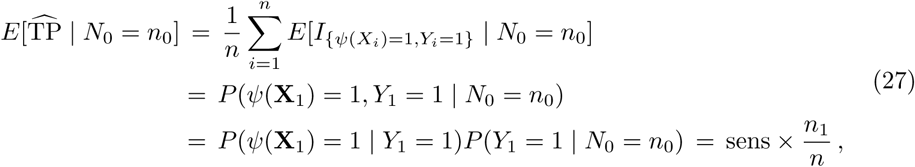

where we used the fact that the event {*ψ*(**X**_1_) = 1} is independent of *N*_0_ given *Y*_1_ and (20). Notice that the equality *P*(*ψ*(**X**_1_) = 1 | *Y*_1_ = 1) = sens depends on the fact that *ψ* is assumed to be constant, so that **(X**_1_, *Y*_1_) behaves as an independent test point (also because of a constant *ψ*, there is no expectation around sens). Hence, 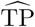 is biased under separate sampling, with

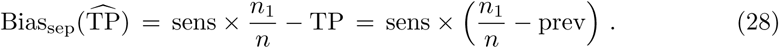

As in the case with the bias of 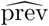 under separate sampling, the bias of 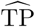 cannot be reduced with increasing sample size. The bias is in fact larger the more sensitive the classifier is. One can derive similar results for 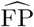, 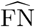, and 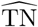.

Perhaps surprisingly, the recall estimator is approximately unbiased under separate sampling:

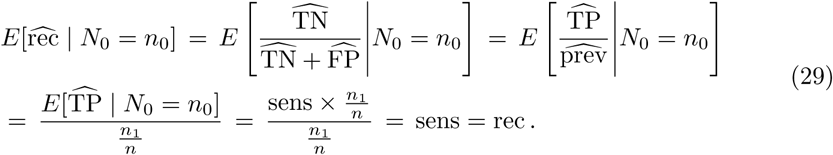

This is a consequence of recall’s not being a function of the prevalence. However, for the precision estimator,

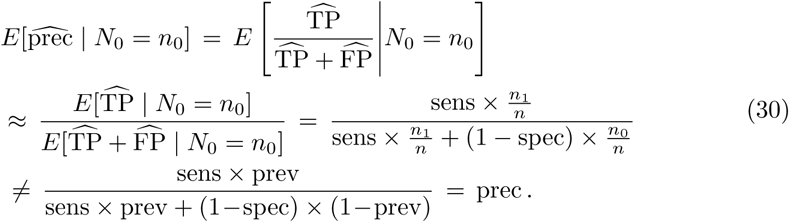

The precision estimator is thus biased under separate sampling, and the bias is larger the further away the true prevalence is from 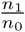. In particular, the bias will be large when prev is small, which is the case in case-control studies involving rare diseases.

### 2.5 Proposed Precision Estimator

In case the true prevalence is known, e.g., from public health records and government databases, then we propose the following estimator of the precision, based on (11):

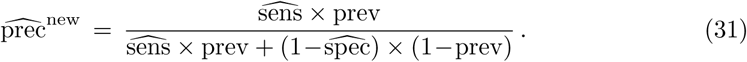

This estimator is still asymptotically unbiased under mixture sampling (which can be seen by repeating the steps in the analysis of the ordinary precision estimator). Under separate sampling, we have

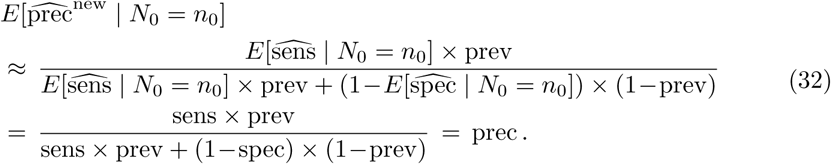

since 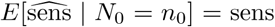 and 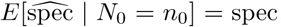, as can be easily shown. Hence, 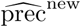 is an asymptotically unbiased estimator of the precision under either mixture or separate sampling. The ordinary precision estimator 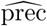 should never be used under separate sampling, or large and irreducible bias may occur. On the other hand, in the impossibility of obtaining information on the true prevalence value, then no meaningful estimator of the precision is possible.

## 3 RESULTS AND DISCUSSION

In this section, we employ synthetic and real-world data to investigate the performance of the proposed precision estimator and the accuracy of the theoretical analysis in Section 2.4. We present results for the bias of the usual and proposed precision estimators under separate sampling. Corresponding results for mixture sampling and the recall estimator can be found in the Supplementary Material.

### 3.1 Experiments with Synthetic Data

We performed a set of experiments employing synthetic models with class-conditional 3-dimensional Gaussian distributions *N*(***μ****_i_*, **∑***_i_*), for *i* = 0, 1, with ***μ***_0_ = (0, 0, 0), ***μ***_1_ = (0, 0, *θ*), where *θ >* 0 is a parameter governing the separation between the classes, and 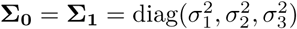 (i.e., a matrix with 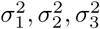 on the diagonal and zeros off diagonal). We consider two sample sizes, *n* = 30 and *n* = 200, so that we can compare the results for small and large sample sizes. All experiments with separate sampling are performed with sample prevalence 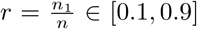, where the value of *n*_1_ is set to *n*_1_ = ⌈*nr*⌉. The synthetic data parameters are summarized in Table 1.

**Table 1.** Synthetic data parameters.

| Parameter | Value |
| --- | --- |
| Dimensionality/ feature size | $D = 3$ |
| Mean difference | $\theta = 2$ |
| Covariance matrix | $\sigma_1^2 = 0.5, \sigma_2^2 = 0.5, \sigma_3^2 = 1$ |
| Sample size | $n = 30, 200$ |
| Sample prevalence $n_1/n$ | $r = 0.1, 0.3, 0.5, 0.7, 0.9$ |
| Actual prevalence | prev = 0.1, 0.3, 0.5, 0.7, 0.9 |

For each value of *r* and prev, we repeat the following process 1,000 times, and average the results to estimate expected error values:

1. Generate sample data *S_n_* of size *n* according to *r* (separate sampling) or prev (mixture sampling);
2. Train a classifier using one of three classification rules [11]: Linear Discriminant Analysis (LDA), 3-Nearest Neighbors (3NN), and a nonlinear Radial-Basis Function Support Vector Machine (RBF-SVM).
3. Obtain recall and precision estimates. Compute both the usual precision estimate **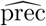** and the proposed **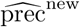**.
4. Obtain (good approximations of) the true precision values, by using a test set of size 10,000.

Fig. 2 displays the results of the experiment. Notice that there is only one curve for the traditional precision estimator **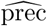** because it does not employ the actual value of prev. The results show that at *n* = 30, all estimators display bias, which is however much larger for the traditional precision estimator. At *n* = 200, the bias of the proposed precision estimator nearly disappears for LDA and is reduced for the other classification rules. Among these classification rules, LDA is the only one with a finite VC dimension, and so the bias in this case is predicted to shrink to zero as sample size increases, according to the theoretical analysis in Section 2.4. Notice also that the bias of the traditional precision estimator is largest when *r* = *n*_1_/*n* is far from prev, and it cannot be reduced by increasing sample size. All these observations are in agreement with the theoretical analysis in Section 2.4 (the results in the Supplementary Material also confirm the theoretical analysis).

**Fig 2.**
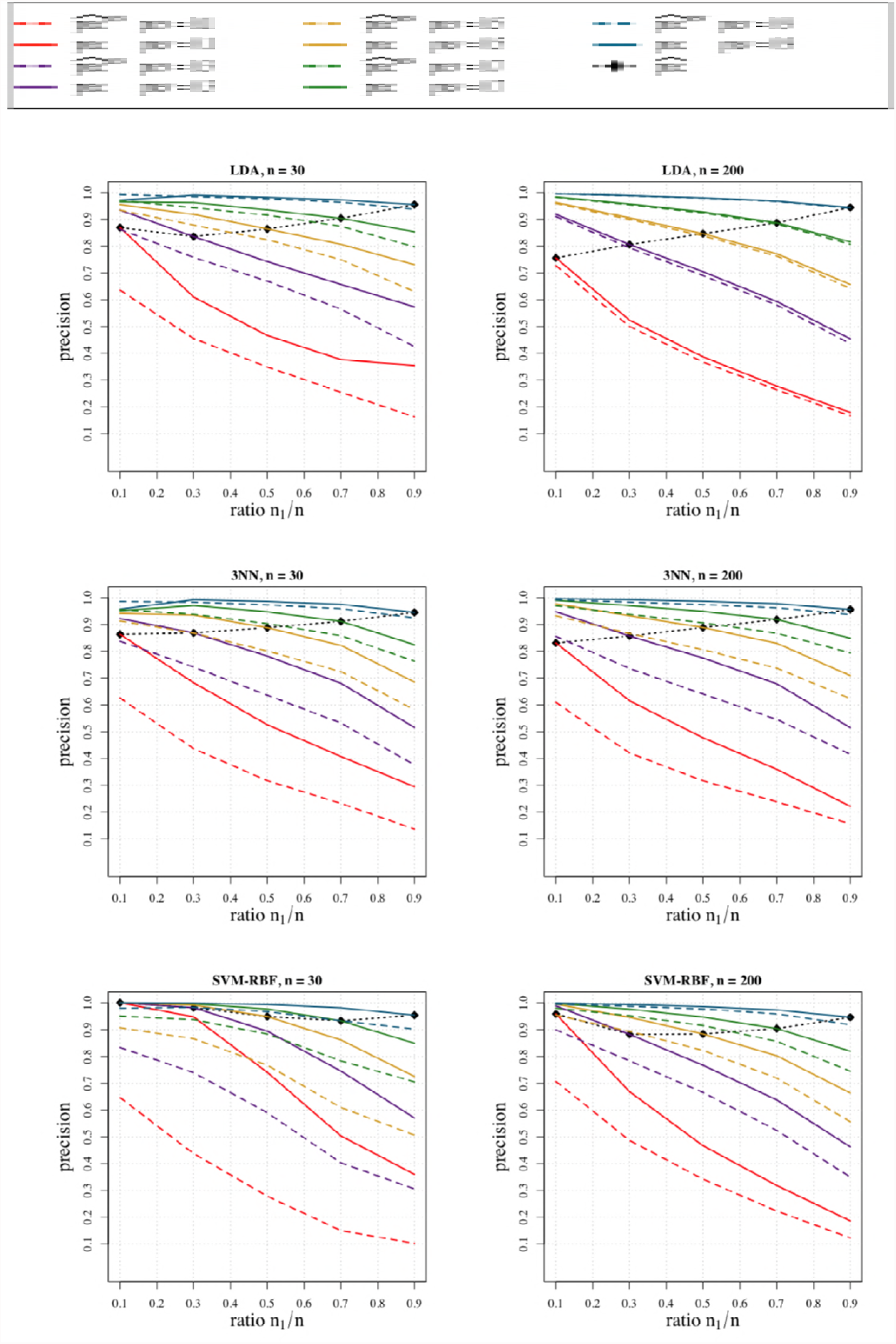
Average true precision values (solid curves) and average precision estimates 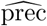 (dash-diamond curve) and 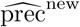 (dashed curves), for LDA, 3NN and RBF-SVM, sample sizes *n* = 30 (top row) and *n* = 200 (bottom row) and different prevalence values as a function of the sample prevalence *r* = *n*_1_/*n*.

### 3.2 Two Case Studies with Real Data

To further investigate the bias of precision estimation under separate bias, we use real data from two published studies. The first [12] uses a tumor microarray dataset containing two types of human acute leukemia: acute myeloid leukemia (AML) and acute lymphoblastic leukemia (ALL). Gene expression measurements were taken from 15,154 genes for 72 tissue specimens, 47 ALL type (class 0) and 25 AML type (class 1), so that the sample prevalence is *r* = 0.347. We computed the traditional the precision estimator 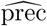, and the proposed estimator 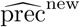 by using the value prev = 0.222, which is the incidence rate of ALL over AML in the U.S. population [13], for four classification rules: Naive Bayes (NB) [14], C4.5 decision tree [15], 3NN and SVM. Fig. 3 displays the results. We can observe that all 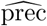 estimates are larger than the more precise 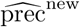 estimates, pointing to an optimistic bias of the usual precision estimator.

**Fig 3.**
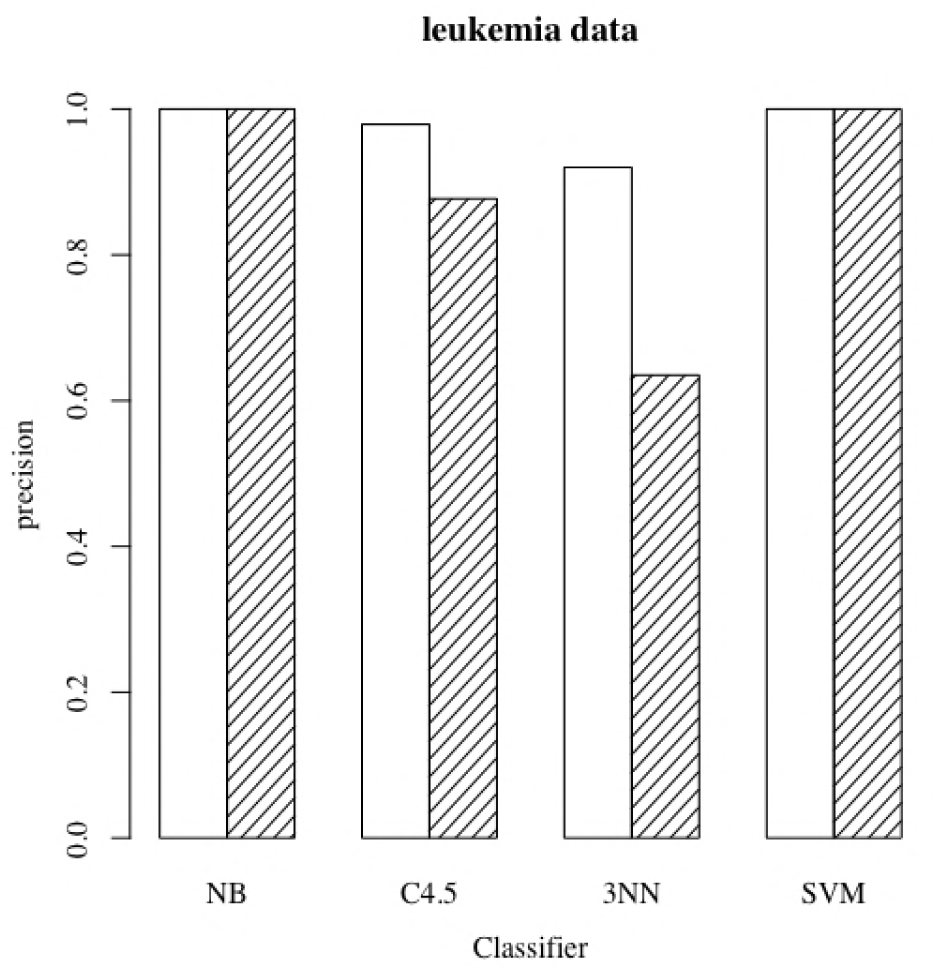
The white bars are the usual estimated precision on the separately-sampled ALL-AML dataset in [12] for four classification rules; the shaded bars are the precision estimates using the prevalence of ALL over AML in the U.S. population.

In the second case study, we employ the liver disease dataset in [16], taken from University of California-Irvine (UCI) Machine Learning Repository [17]. This data set contains 5 blood tests attributes and 345 records in which 145 belong to individuals with liver disease (class 0) and 200 measurements are taken from healthy individuals (class 1), so that *r* = 0.42. The authors in [16] constructed classifiers for diagnosis of liver disease based on the blood tests variables in this data set, and reported the estimated accuracy, precision, sensitivity and specificity for five classification rules: Naive Bayes (NB), C4.5, 3NN, Back-Propagated Neural Network [11] and a Linear SVM. This dataset was donated to UCI in 1990, and according to [18], the prevalence rate for chronic liver diseases in the US was prev = 0.1178 between 1988 and 1994, which we use as the approximated prevalence in the computation of the 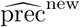 estimator. Fig. 4 displays the results. As in the previous study, we observe that all 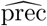 are larger than the more precise 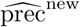 estimates, but this time the difference is much larger, indicating possible strong optimistic bias of the traditional precision estimator.

**Fig 4.**
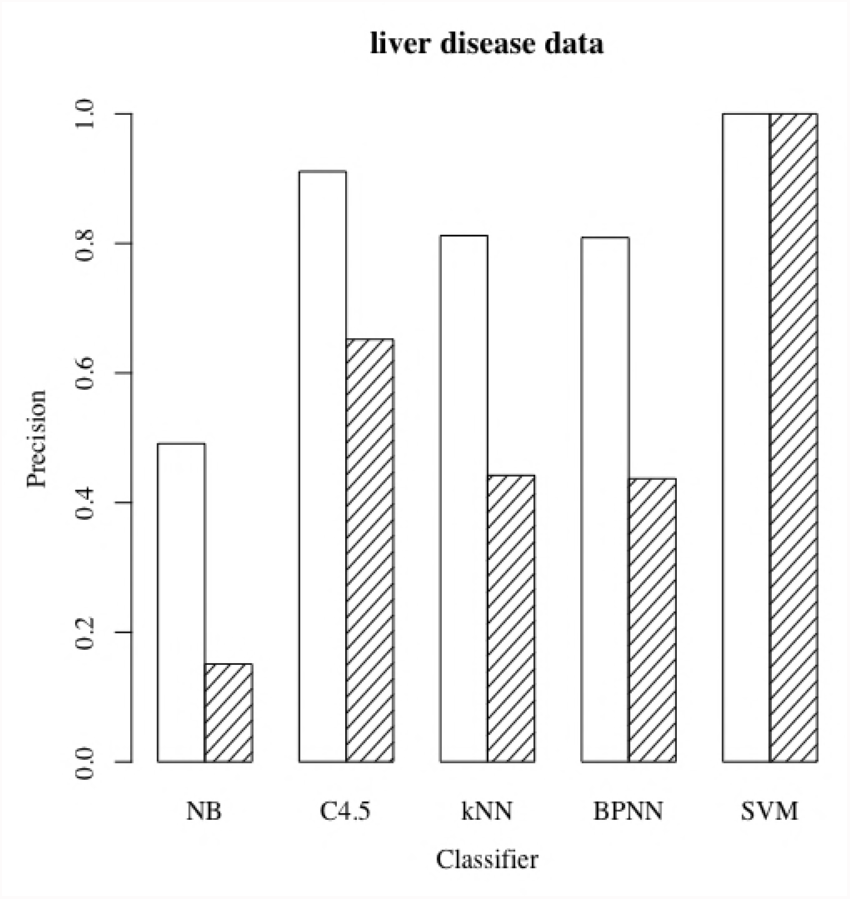
The white bars are the usual estimated precision on the separately-sampled liver disease dataset in [16] for four classification rules; the shaded bars are the precision estimates using the prevalence of liver disease in the U.S. population.

## 4 Concluding Remarks

Accuracy and reproducibility in observational studies is critical to the progress of biomedicine, in particular, in the discovery of reliable biomarkers for disease diagnosis and prognosis. In this study, we showed, using analytical and numerical methods, that the usual estimator of precision can be severely biased under the typical separate sampling scenario in observational case-control studies. This will be true especially in the case of rare diseases, when the true disease prevalence will be small and differ significantly from the apparent prevalence in the data. If knowledge of the true disease prevalence is available, or can even be approximately ascertained, then it can be used to define a new precision estimator proposed here, which is nearly unbiased at moderate sample sizes. Absence of knowledge about the true prevalence means simply that the precision cannot be reliably estimated in observational case-control studies.

## Acknowledgments

The authors acknowledge the support of the National Science Foundation, through NSF award CCF-1718924.

